# Whole genome sequencing elucidates the species-wide diversity and evolution of fungicide resistance in the early blight pathogen *Alternaria solani*

**DOI:** 10.1101/2021.06.28.450143

**Authors:** Severin Einspanier, Tamara Susanto, Nicole Metz, Pieter J. Wolters, Vivianne G.A.A. Vleeshouwers, Åsa Lankinen, Erland Liljeroth, Sofie Landschoot, Žarko Ivanović, Ralph Hückelhoven, Hans Hausladen, Remco Stam

## Abstract

Early blight of potato is caused by the fungal pathogen *Alternaria solani* and is an increasing problem worldwide. The primary strategy to control the disease is applying fungicides such as succinate dehydrogenase inhibitors (SDHI). SDHI-resistant strains, showing reduced sensitivity to treatments, appeared in Germany in 2013, five years after the introduction of SDHIs. Two primary mutations in the Sdh complex (SdhB-H278Y and SdhC-H134R) have been frequently found throughout Europe. How these resistances arose and spread, and whether they are linked to other genomic features, remains unknown.

We performed whole-genome sequencing for *A. solani* isolates from potato fields across Europe (Germany, Sweden, Belgium, and Serbia) to better understand the pathogen’s genetic diversity in general and understand the development and spread of the genetic mutations that lead to SDHI resistance. We used ancestry analysis and phylogenetics to determine the genetic background of 48 isolates. The isolates can be grouped into 7 genotypes. These genotypes do not show a geographical pattern but appear spread throughout Europe. The *Sdh* mutations appear in different genetic backgrounds, suggesting they arose independently, and the observed admixtures might indicate a higher adaptive potential in the fungus than previously thought.

Our research gives insights into the genetic diversity of *A. solani* on a genome level. The mixed occurrence of different genotypes and apparent admixture in the populations indicate higher genomic complexity than anticipated. The conclusion that SDHI tolerance arose multiple times independently has important implications for future fungicide resistance management strategies. These should not solely focus on preventing the spread of isolates between locations but also on limiting population size and the selective pressure posed by fungicides in a given field to avoid the rise of new mutations in other genetic backgrounds.

## Introduction

Early blight and Late blight are the two main diseases that can cause severe damage to European potato production. The causal agent of early blight is the fungal pathogen *Alternaria solani*. If the weather conditions are favorable and *A. solani* remains uncontrolled, yield losses can reach up to 40% (Leiminger & Hausladen, 2012, 2014). Typical symptoms of potato early blight are dark brown to black lesions with concentric rings. In the beginning, the necrotic area is often surrounded by a yellow, chlorotic halo (Rotem, 1994). Due to climatic changes Delgado-Baquerizo *et al*., (2020), predicted a global increase of soil-borne pathogens, e.g., *Alternaria* spp., which would imply increasing disease pressure in the field. The potato crop is the fourth most important field crop in European agriculture based on yield (tonnes/year), after wheat, maize, and barley (Eurostat 2020). To guard the yield, farmers adhere to good agricultural practices, e.g., crop rotation and sufficient nutrient supply, but in most cases the application of fungicides in needed. For early blight control, three main fungicide groups are available: the Quinone outside inhibitors (QoIs), the Succinate Dehydrogenase Inhibitors (SDHIs) and the Demethylation inhibitors (DMIs), of which QoIs and SDHIs are most widely used and until recently considered most effective.

The development of resistance has been reported in *A. solani* and other fungi for all of these fungicides groups. DMI resistance appears to be related to changes in expression of the target site, Cyp51, possibly linked with a mutation (Zhang *et al*., 2020). Mutations in *Cyp51* genes are common in other pathogens to which DMIs are applied (Pereira *et al*., 2017; Blake *et al*., 2018). For the other two fungicide groups, mutations in the pathogen target sites have already been confirmed in *A. solani*. The F129L mutation in the cytochrome b of the mitochondrial electron transport complex III (Bartlett *et al*., 2002) frequently occurs in the field and reduces the efficiency of QoI-fungicides to a certain degree (Leiminger *et al*., 2014; Odilbekov *et al*., 2016). Concerning the SDHIs, several different mutations have been identified in the subunits b, c, and d of the *sdh*-gene in the mitochondrial electron transport complex II: *Sdh*B-H278Y, *Sdh*B-H278R, *Sdh*C-H134R, *Sdh*C-H134Q, *Sdh*D-D123E, *Sdh*D-H133R (Mallik *et al*., 2014; Metz *et al*., 2019). For these SDH mutations, a negative impact on fungicide efficiency has also been shown in several studies (Landschoot *et al*., 2017; Metz *et al*., 2019).

In a European study, 70% of isolates sampled between 2014 and 2015 contained one of the SDHI mutations and 40% also had the cytochrome b F129L mutation, leading to QoI resistance and thus possessed a dual fungicide resistance (Landschoot *et al*., 2017). A more recent, long-term Swedish field study, showed that nearly all Swedish isolates now carry the F129L mutation (Edin *et al*., 2019). In Germany also nearly all samples have the F129L mutation and 43% show mutations in SDH subunits (Nottensteiner *et al*., 2019). In the United States, a study of over 1000 isolates revealed that between 2013 and 2015 the presence of the F129L mutation rose from 92 to 99% and mutations in any of the SDH subunits were also present in 99% of the isolates (Bauske *et al*., 2017).

The genetic diversity of *A. solani* in the field is relatively understudied. Possibly, because *A solani* historically has been described as an asexual species and limited variation was expected. Yet, considerable genetic diversity was found in South African isolates using random amplified microsatellite (RAMS) primers (van der Waals *et al*., 2004). Random amplification of polymorphic DNA (RAPD) profiling confirmed this surprisingly high genetic diversity of field isolates in Germany (Leiminger *et al*., 2016) or Wisconsin (Weber & Halterman, 2012). Also on tomato in India, a marker-based study revealed higher within and between state diversity in *A. solani* than anticipated by the authors (Upadhyay *et al*., 2019). Barcode sequencing revealed the presence of multiple *A. solani* haplotypes in a single field (Adhikari *et al*., 2020).

Odilbekov found that genetic composition of the pathogen populations in the field changed after fungicide treatment. Their study revealed an increase of genotypes with reduced sensitivity over time (Odilbekov *et al*., 2019). However, the use of Amplified Fragment Length Polymorphisms (AFLP) limited the resolution of the genetic diversity analysis and only two main genotypes could be described. Genome-wide sequencing approaches such as Genotyping By Sequencing (GBS) or full genome sequence analyses, are very powerful tools that are able to reveal genomic variation that previously remained hidden (Everhart *et al*., 2020). Indeed, a GBS study revealed that microevolutionary factors might play an important role in population structure of various *Alternaria spp* (Adhikari *et al*., 2019).

Whereas fungicide resistance is on the rise in *A. solani*, it is not clear whether the causal mutations occur in one or few genetic backgrounds and spread or whether they arose multiple times independently in different genetic backgrounds. Understanding such microevolutionary forces at play in and between populations and their roles in fungicide resistance evolution is important. If mutations arise only once and then spread, this should have implications for management practices, for instance on transportation and movement of tubers and infected material.

Here we combine a genomic diversity study with analyses of fungicide resistance targets. We make use of the recently published *A. solani* reference genome (Wolters *et al*., 2018) and a Europe-wide selection of *A. solani* samples to study the genetic diversity of the isolates in general and the occurrence of SDHI resistance in a genomic context.

## Methods

### Fungal Material

48 different isolates were collected from five different localities across Europe (Figure 1). Isolates DE_NM014 – DE_NM020 were collected from potato fields in Bavaria, Germany, DE_NM006 – DE_NM013 from fields in Lower Saxony in Germany. SE_EL001 – SE_EL012 isolates originate from Southern Sweden, BE_SL001 – BE_SL008 from fields in Belgium, and RS_ZI001 – RS_ZI008 were collected from two localities in Serbia. US_JW001 – US_JW004 and US_NM022 were collected in the United States of America. A detailed overview can be found in Table 1. All isolates were samples in their respective fields from symptomatic potato leaves. Leaves were dried between tissue paper and surface sterilized I before placement on SNA (0.2 g/l glucose, 0.2 g/l sucrose, 0.5 g/l MgSO_4_-7H_2_O, 0.5 g/l KCl, 1.0 g/l KH_2_PO_4_, 1.0 g/l KNO_3_, 22.0 g/l agar; 600 µl/l 1M NaOH). A single spore was collected from each isolate and propagated under sterile condition for further usage.

**Figure 1.**
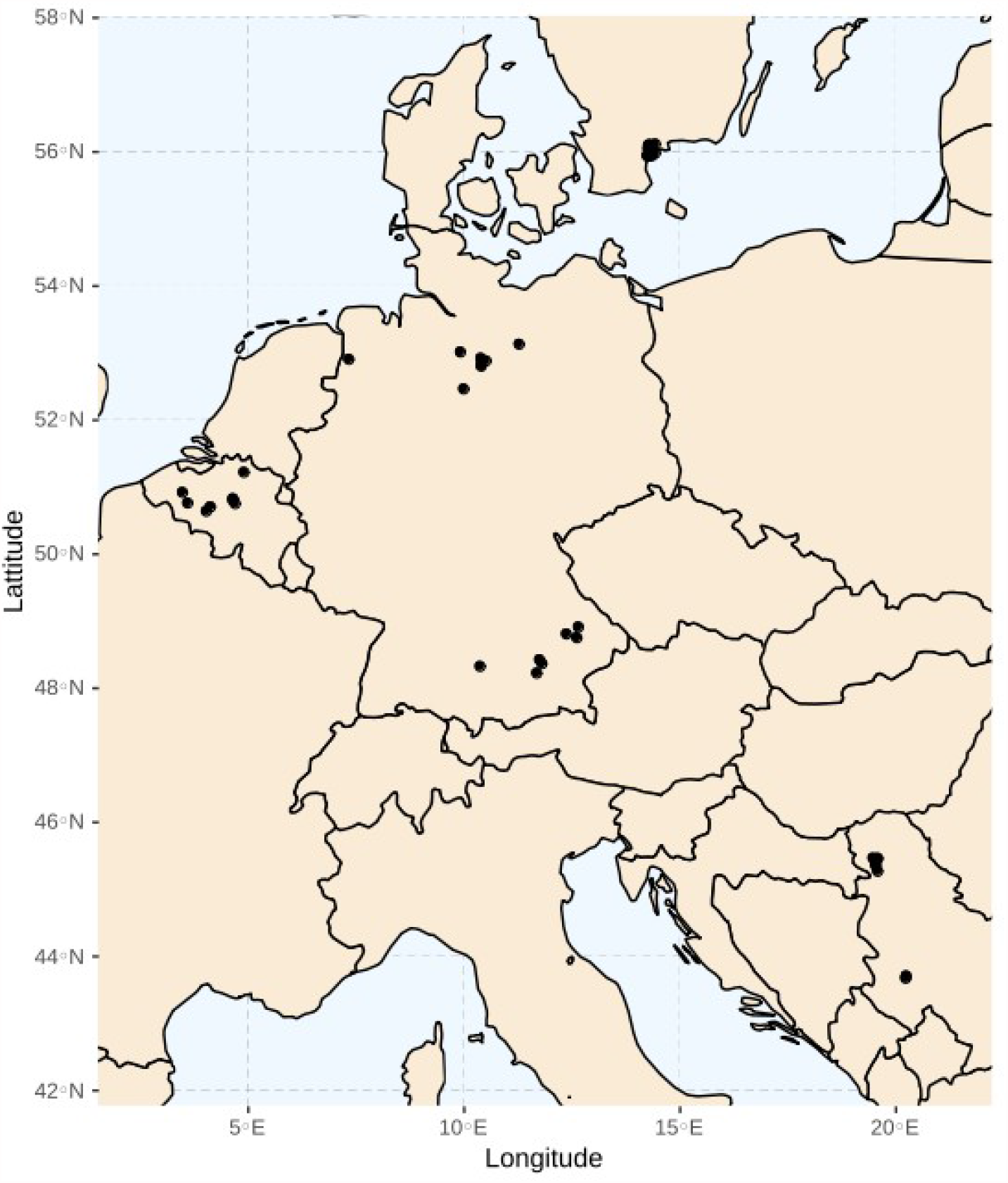
Overview map of the sampling locations of the A solani isolates used in this study.

### DNA extraction

For high quality DNA extraction, 200ml of potato dextrose broth was inoculated with small agar plugs from 10 to 14 day old *A. solani* cultures, grown on SNA. Liquid cultures were incubated for 48 hours at 28°C on a rotary shaker (110rpm). Afterwards, mycelium was filtered through a cheese cloth and squeezed to remove most of the liquid. After freeze drying, mycelium was grinded with liquid nitrogen and a bit of clean sea sand to break the cells most efficiently. 50-100 mg of lyophilized powder was added to 1ml of DNA extraction buffer (containing extraction buffer (0.35M sorbitol, 0.1M Tris-HCl pH7.5, 5mM EDTA), nucleic lysis buffer (2% CTAB, 2M NaCl, 0.2M Tris-HCl pH7.5, 50mM EDTA) and sarkosyl (10%) in a ratio of 2.5:2.5:1), 20µl proteinase K (25mg/ml) and 20µl RNAse A (20mg/ml) in a 2ml tube. Everything was mixed well and incubated for 1hour at 60°C – invert the tubes occasionally to gently mix the content. Afterwards samples were centrifuged for 10 minutes at maximum speed and supernatant was transferred into a new tube. An equal amount of PCI (Phenol:Chloroform:Isoamylalcohol (24:24:1)) was added to the tube and incubated overnight on a rotary shaker at 4°C (gentle rotation to avoid damaging DNA). On the following day, samples were spun down for 15 minutes at maximum speed and upper phase was transferred to a new tube. An equal amount of SEVAG (Chloroform:Isoamylalcohol (24:1)) was added and tubes were incubated for at least 1 hour on a rotary shaker at 4°C (gentle rotation). In a next step, samples were centrifuged for 10 minutes at maximum speed and supernatant was transferred to a new tube. For precipitation, 0.7 volume of isopropanol (room temperature) was added and samples were spun down for 15 minutes at 10.000 rpm. After washing the DNA pellet with 70% ethanol, remaining liquid was removed by pipetting. To entirely remove the ethanol, tubes were placed in a heating block at 37°C to let them dry completely. DNA pellet was dissolved in 100µl distilled water and stored at 4°C. A Qubit was used to determine the quantity of the extracted DNA.

### Mapping and SNP processing

The quality of the raw sequence data was checked using FastQC. The Burrows-Wheeler alignment tool (Li & Durbin, 2009) was used to map all samples to our reference genome NL03003 (Wolters *et al*., 2018). Reads deduplication was performed using the picard tool Markduplicate. Subsequently, the single-nucleotide polymorphism (SNP) was called using GATKs Haplotype-Caller in GVCF mode (McKenna *et al*., 2010) with default settings, except for having ploidy set to one. The SNPs were filtered using GATK with modified parameters compared to the recommended filtering criteria. SNPs were filtered out when they met the following parameters: low mapping quality rank sum test (MQRankSum < - 12.5), low quality by depth (QD < 2.0), low read pos rank sum test (ReadPosRankSum < −8.0), high Fisher strand difference (FS > 60.0), low RMS mapping quality (MQ < 40.0), high strand odds ratio (SOR > 3.0), high haplotype score (HaplotypeScore > 13.0). Additionally, the insertion or deletions (indels) were filtered out using GATK with the following parameters: low quality depth (QD < 2.0), low read pos rank sum test (ReadPosRankSum < -20.0), and high Fisher strand difference (FS > 200.0). Afterwards, SNP clusters and SNPs close to indels were removed using SnpSift filter (Cingolani *et al*., 2012a). The summary of the variant analyses were generated by SnpEff (Cingolani *et al*., 2012b).

### Visual Inspection

The vcfR package was used to read and visualize the vcf file for quality control (Knaus & Grünwald, 2017). The read depth (DP), the mapping quality (MQ), the Phred-scaled quality (QUAL), and the variants were visualized. This identified several low confidence areas with very high DP. The masker() function was used to filter out data outside the confidence range: DP (0-2500) and MQ (50-60).

### Phylogeny of *A. solani* samples and extraction of SDHI gene data

In order to validate the nature of our samples, we extracted three commonly used barcode genes *gapdh, rbp2* and *tef1*, that can be used to distinguish *Alternaria* spp (Woudenberg *et al*., 2013). References for *gapdh* (KC584139), *rbp2* (KC584430) and *tef1* (KC584688) were extracted form NCBI search against the reference genome using BLAST. Start and Stop coordinates were extracted and alternative reference fasta sequences were extracted using GATK (McKenna *et al*., 2010). All sequences were loaded in ab12phylo (Kaindl *et al*., 2021), and a multi-gene phylogeny was constructed using built-in RaxML-NG (Kozlov *et al*., 2019) including *A. solani* reference sequences and the sequences of two closely related sister species *A. dauci* and *A. porri* (GAPDH (KC584111, KC584132), RBP2 (KC584392, KC584421), TEF1 (KC584651, KC584679) for *A. dauci* and *A*.*porri* respectively). To analyze the genomic regions coding for the SDH subunits, we performed a BLAST search of sequences each of the SDH subunits against the *A. solani* reference and used the identified start and stop site to extract alternative reference fasta files using GATK (McKenna *et al*., 2010).

### Genetic diversity summary statistics

Basic genomic statistics were calculated with the R package PopGenome (Pfeifer *et al*., 2014). For whole genome-analysis, vcf files were split into smaller chunks using vcftools (Danecek *et al*., 2011) and imported using the readData function(include.unknown=F, format=“VCF”, SNP.DATA=T, big.data=T). Statistics were generated via neutrality.stats() and diversity.stats() and for further use transformed into R-dataframes. Populations were defined via the set-populations() function. The Site Frequency Spectrum (SFS) was generated by extraction of AC-values from the vcf file. This was done using the commands strsplit(), do.call() and table() in base R. Basic histograms were plotted using ggplot2 (Wickham, 2009).

Sliding window analysis of SNPs per site was performed using the sliding.window.transform() option of PopGenome (w=10000, j=10000, type=2, whole. data=T). To achieve that, vcf chunks of chromosomes were imported separately and analyzed discretely. Therefore, a function was defined to allow the analysis of each chromosome and to re-aggregate generated data, subsequently. This is necessary since the native function for data-import (readVCF) does not apply for haploid datasets. Segregating sites per window were extracted and normalized per site. ggplot2 was used to visualize our findings.

### Population phylogeny

We extracted the SNPs as the alternative reference genomes using the GATK FastaAlternateReferenceMaker tool to create an alignment of the whole isolates’ genomes and to construct a population phylogeny (DePristo *et al*., 2011). To create a maximum likelihood phylogeny RAxML version 8 (Stamatakis, 2014) was used with -p 1122590 -f a -x 1122590 -m GTRGAMMA -# 100 -s input -n output and 100 bootstraps. The phylogenetic tree was vizualized using the R packages ggtree and treeio (Wang *et al*., 2020).

### PCA (Principal Component Analysis)

As a dimensionality-reduction technique, principal component analysis (PCA) was generated using the R packages gdsfmt and SNPRelate (Zheng *et al*., 2012) and the PCA was visualized using the ggplot2. To avoid the overlapping labels on the PCA, the R package ggrepel was used (Slowikowski, 2018).

### LEA

Ancestry analysis was performed using the R package LEA (Frichot & François, 2015). For that, vcf files were converted into a genotypic matrix using vcftools (--plink) and the LEA-function “ped2geno”. This file was then transformed manually (9=1, 2=0). In the LEA package, *sparse nonnegative matrix factorization* (sNMF) was performed. A cross-entropy criterion was calculated to identify the best statistical model describing ancestral populations of the dataset. The minimal cross-entropy for the dataset was determined with multiple repetitions (K1:15, ploidy=1, entropy=T, rep=10). Admixture analysis was performed for k=6, k=7 and k=8 [*Q(obj, k, run=which*.*min(cross*.*enropy(obj, k)))*] and ordered according to individually assigned genotypes or sample origin. Visualization and inference with mutations was done using tidyverse package (Wickham, 2019) and ggplot2.

### Distance matrix

The SNP distance matrix was constructed from the alternative reference sequences with the extract alternative reference fasta option from GATK (McKenna *et al*., 2010). Two versions were created. One including all sites, one removing all incomplete cases (e.g. removing sites that were called in one isolate, but not in another). The matrix was created using snp-dists -a (https://github.com/tseemann/snp-dists).

### Data availability

Raw short read data are deposited to NCBI SRA. SNP call files are deposited to Zenodo.org under 10.5281/zenodo.5029386.

## Results

### *A. solani* is highly polymorphic across Europe

To assess the genome-wide diversity of *A. solani* in Europe, we collected 43 isolates from potato fields in Belgium, Northern Germany (Lower Saxony), Southern Germany (Bavaria), Serbia, and Sweden. The sample numbers are roughly equal between locations and represent a cross section of Europe (Figure 1, Table 1). We also include 5 isolates originating from the United States to allow comparisons. All isolates were sequenced using Illumina HiSeq and had an average read depth of 62x coverage, after mapping and filtering (Table 1). On average we found 47,098 SNPs in a single isolate, when comparing to the reference, corresponding to one SNP every ∼700 bases. The number of SNPs in most isolates ranged from 32,079 SNPs (in DE_NM019) to 52,911 (in DE_NM017). BE_SL002 appeared to be an outlier with 157,798 SNPs compared to the reference. However, extraction of the sequences of typical barcode markers used in phylogenetic analyses of *Alternaria* species (RPB2, GAPDH and EF1) confirmed that BE_SL002 is *A. solani*. It belongs to one of the two major haplotypes that can be detected in our dataset based on these barcodes and does not cluster with one of the related sister species (Figure S1).

In total we find 262,928 (136,180 without BE_SL002) SNPs between all isolates. Due to very the large number of unique SNPs in BE_SL002, compared to all other samples, further analyses were done without this sample, to allow better comparison.

We find a transition vs transversion (ts/tv) ratio of 3.05. Forty-eight percent of SNP are singletons. The Site Frequency for the sample set shows a homogenic decrease from singletons to doubletons. However, the number of SNPs is not decreasing continuously, in fact, some minor peaks are occurring for higher SNP frequencies (Figure S2). This indicates that there are no large population expansions (as would be the case with increased number of singletons). The slightly uneven distribution might be an effect of partial clonality in the sample set. Next, we calculated several population genetics summary statistics for the sample set. Overall, Watterson θ per site equals 0.019 and π equals 0.016. The overall Tajimas’ D of the dataset is rather neutral with a value of 0.629.

On average we find one SNP every ∼241 bp. SNP density varies per chromosome, ranging from on every 197 bp for chromosome 4 to one every 277 bp for chromosome 2. Using sliding window analysis with 10kb windows, confirms that SNP-density varies along the chromosomes and is highest and lowest in chromosome 6 and 1 respectively (Figure S4). The different genes coding for the different *Sdh* subunits that are prone to fungicide resistance mutations are located on chromosomes 7 (*SdhB*), 2 (*SdhC*) and 5 (*SdhD*). Thus, the fungicide target genes are not located specifically on very conserved or very diverse chromosomes.

### *A. solani* shows differences in diversity between locations

Next, we wanted to know whether the genetic variation is similar for each of the five samples locations in Europe, and in our US samples. To this end, we summarized all statistics per location (Table 2). Even excluding BE_SL002, the Belgian samples had the highest number of SNPs (103,971 SNPs in total), followed by Bavaria (101,623). Lower Saxony and Serbia have 62,378 and 63,997 SNPs respectively.

Interestingly, in Sweden, from where we analyzed 12 samples, rather than 7 or 8, the number of segregating sites is only 71,232. This might in part be due to the fact that the Swedish samples were all collected in one locality, but it should be noted that with the exception for 1 sample this was also the case for Serbia. Moreover, a clear link between the number of segregating sites and the geographical spread between the samples of a certain locality cannot be made. To illustrate, the furthest distance between samples in Belgium is 107 km, the maximum distance between samples in Lower Saxony is 262 km, yet Belgian isolates remain more diverse.

In line with the higher number of SNPs, we also find that the Belgian samples have the highest nucleotide diversity (Watterson’s θ or π). Interestingly, we found some variation in Tajima’s D. However, Tajima’s D is negative or close to 0 in most localities. This indicates a relatively high number of singletons and is in line with the overall observations from the Site Frequency Spectrum (SFS). However, a Tajima’s D of 1.024 in Sweden seems to violate the assumption of neutral theory suggesting less low-frequency polymorphisms, probably due to higher clonality in these samples.

### Classification of *A. solani* genotypes

We hypothesized that there is variation in genetic diversity and the genetic background of the isolates between each locality. Therefore, we wanted to define the overall population structure and see whether genotype distribution of *A. solani* matches with respective sample origin, e.g. genotypes showing regional clustering. We first reduced the complexity of the data by constructing a PCA for the SNPs (Figure 2A). Combined the first two components explain 37% of the variation. When looking at the PCA, one can identify three or four clusters, each of varying size and spread. All clusters that can possibly be identified contain isolates from different geographical origins. Note that sample BE_SL002 is only on the edge of the PCA for component 2 and not a clear outlier. To further corroborate these findings, we set to construct a phylogenetic tree based on all identified SNPs in the isolates (Figure 2B). As expected, isolates that grouped close together in the PCA also group in the phylogenetic tree. These findings are supported by high bootstrap values. In the phylogenetic tree, BE_SL002 groups together with the isolates that are closest in the PCA, yet it has a longer branch length. Color coding by locality in the PCA and on the branches of the tree shows that most localities indeed harbor several unrelated isolates.

**Figure 2.**
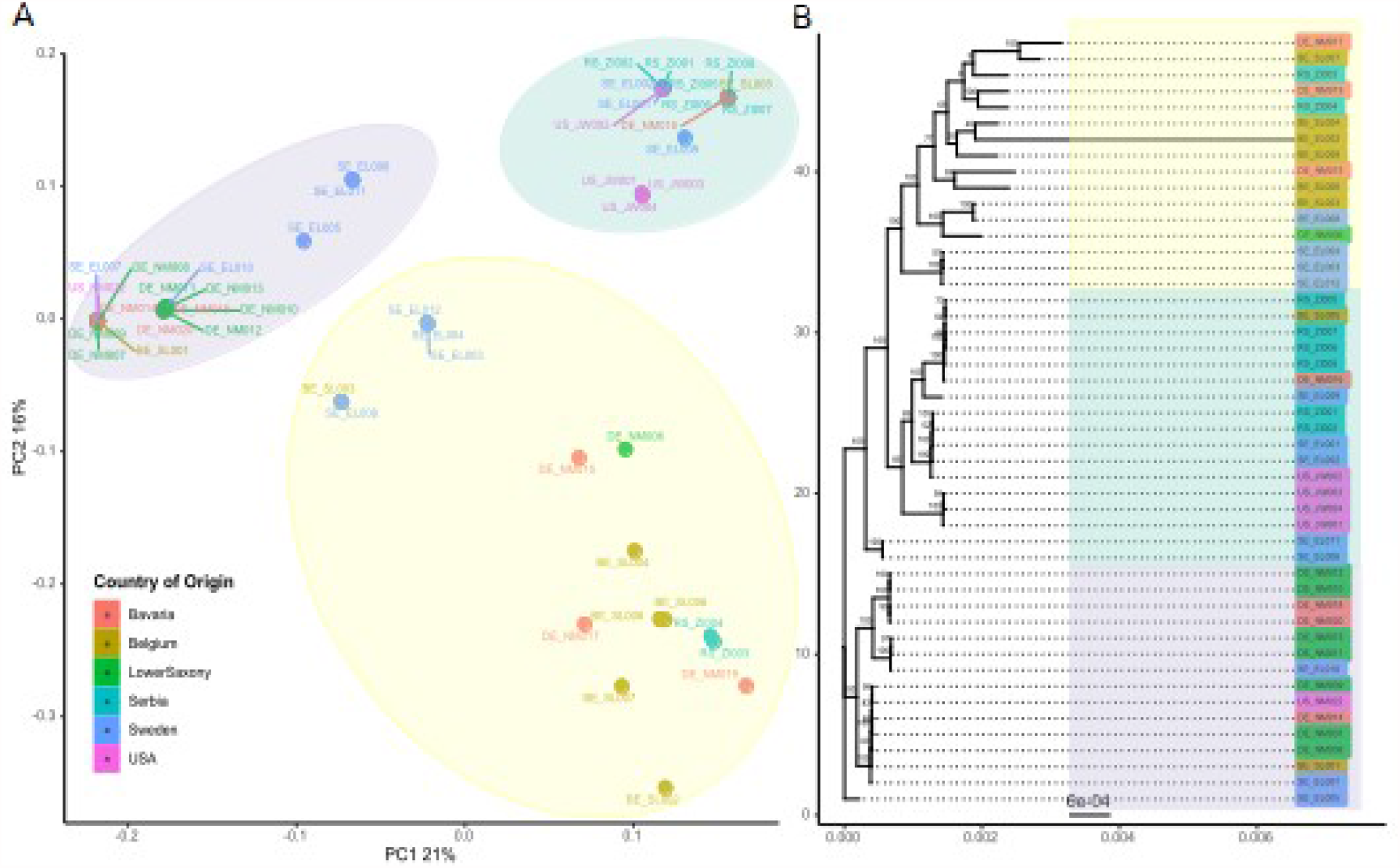
Principal component analysis (PCA) and phylogenetic tree constructed of the 48 *Alternaria solani* isolates. A) Scatter plots of the first two principal components made using SNPRelate and ggplot2 packages in R. The x- and y-axis represents the PC1 (with variance explained 21%) and PC2 (with variance explain 16%). The isolates are color-coded according to their region of origin. Highlighted clusters in yellow, green, and purple are based on the clustering on the phylogenetic tree. B) Phylogeny of reconstructed whole-genome sequences for each of the 48 isolates, made using RAxML (GTRGAMMA) with 100 bootstraps. The phylogenetic tree is constructed using treeio and ggtree packages in R. The color coding of the isolates corresponds to the region of origin.

To further define the population structure of the isolates, we calculated admixture using the R package LEA (excluding BE_SL002). To estimate the most likely number of ancestral populations of the dataset, the cross-entropy criterion was calculated. Lowest cross-entropy can be observed at K=7 (Figure S3), thus admixture analysis was conducted for K = 7 populations as well at K=6 and K=8. With K = 7, we can see groupings similar to those observed in the PCA. Several clusters show mainly uniform, un-admixed ancestry, like the one with BE_SL005, RS_ZI008, RSZI006 and DE_NM016 (all yellow in Figure 3B) also cluster closely together in the PCA and form a monophyletic clade in the phylogeny. All genotypes are present in distinguishable groups. When comparing K = 6 and K = 7, we observe one cluster (US_JW001, US_JW003, US_WJ004) dividing into a clear not-admixed genotypes. In some samples, increasing K can be associated with increasing admixture of various genotypes (e.g. in DE_NM006). For K=7, 13 isolates show a dominant ancestor only with minimal admixture. 18 isolates show admixture of two ancestors and 16 samples show admixture with 3-4 ancestors. Admixture from 5 ancestors can only be found in RS_ZI004.

**Figure 3.**
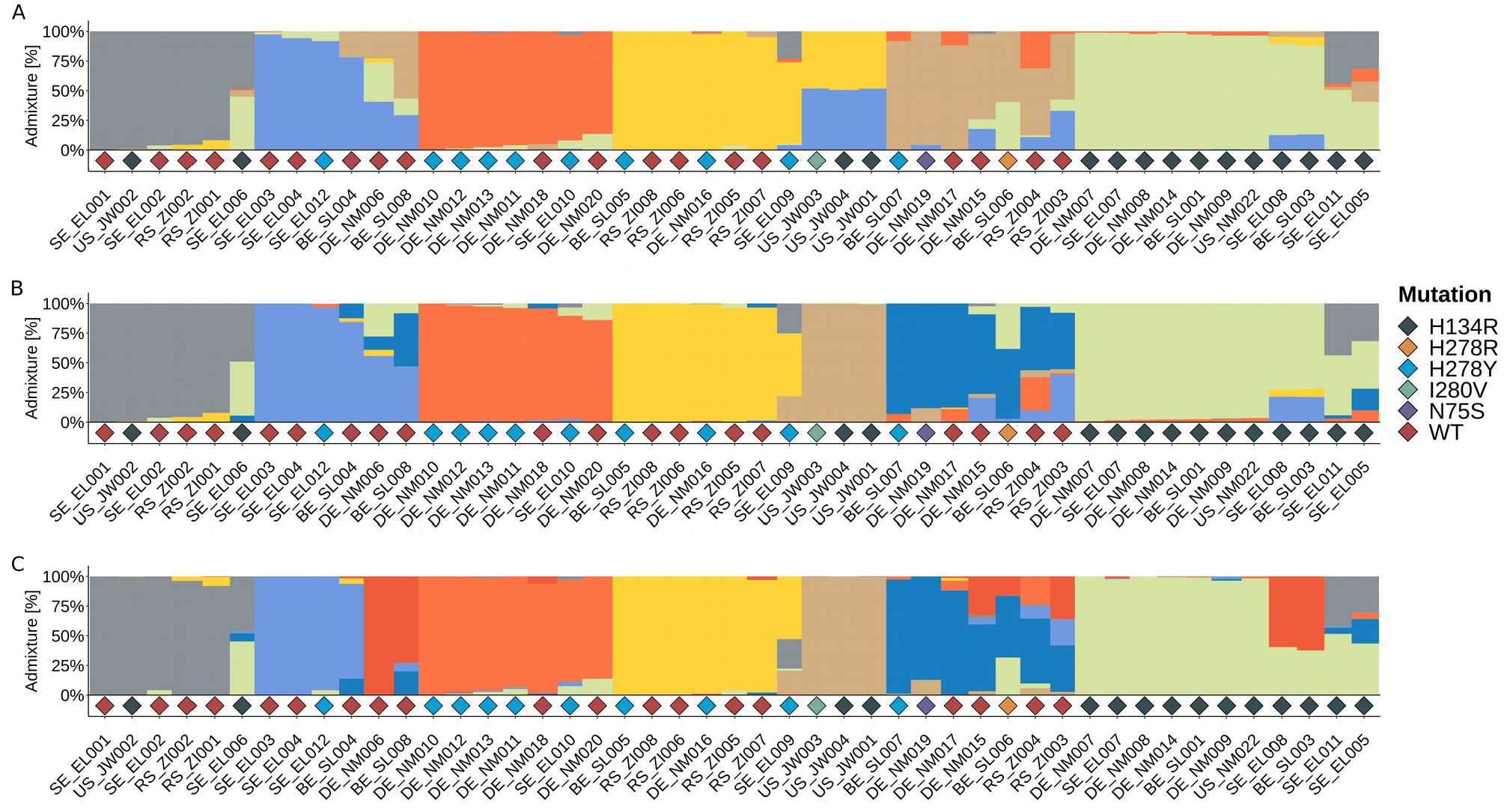
Ancestry analysis on A. solani samples shows both, clear genotypes and highly mixed isolates and no link to SDHI-mutations. Ancestry analysis was performed *A. solani* isolates (n=47, BE_SL002 excluded) using the R-package LEA [*snmf(K=1:15, rep=10, ploidy=1)*] for 6 (A), 7 (B) and 8 (C) ancestral populations. Colors represent individual genotypes assigned by *sparse nonnegative matrix factorization*. Bar plots show the admixture of each genotype in every isolate. Diamonds represent mutations against SDHIs found in the data set.

Next, we assigned each isolate to the respectively most dominant of the 7 genotype groups and calculated diversity statistics for each of them (Table 3). The nucleotide diversity differs within each genotype group. Genotype 6 (dark blue) has the highest numbers of SNPs, 96,239, probably because it is also the most admixed. Interestingly, the number of SNPs of the genotypes does not scale with sample size e.g., genotype 3 (n=7) has only 14,459 SNPs. Therefore, the nucleotide diversity per site diverges strongly between the genotypes, ranging from 0.002 (genotype 4) to 0.023 in genotype 6. Tajimas’ D is ranging from -1.709 in genotype 4 up to 2.212 in genotype 3. Genotypes 2, 6, and 7 show neutral Tajimas’ D, the calculation for Genotype 5 is not possible due to sample size.

Seeing the limited diversity in some genotype groups, we asked how many isolates could be considered clones of each other. To this extend we created a pairwise snp-distance table for isolates (Table S1). This reveals that on average there are 40,282 SNPs between the isolates (35,365 when taking only complete cases (e.g. sites that have been called in all isolates reliably). The lowest number of SNPs between two isolates is 822 (135 with complete cases). Pairwise comparisons with such low number of differences suggest the existence of true clonal isolates. Looking in the genotype groups constructed by LEA, we see that 4 genotype groups contain 3-4 samples that have less than 1,500 SNPs between them and one genotype group shows 2,122 SNPS shared between 6 of its samples. Thus, our Europe-wide sample set consists of a considerable number of likely true clones.

To better visualize the diversity per location, we also sorted the isolates per locality (Figure 4). It should be noted that each location contains a mix of different genotypes. The dominating genotypes of a locality vary slightly, but it is not possible to assign a single dominant genotype to a single region, thus, falsifying our hypothesis that *A. solani* genotypes would show regional clustering. Variation in the diversity within geographical regions can also be observed. The samples from Belgium and Sweden show the most complex admixture structure, followed by Bavaria. The high complexity in Sweden is particularly interesting, because in absolute number of segregating sites, this population did not appear to be the most diverse. Interestingly, some of the clones described above, can be found on different sides of the European continent (e.g. BE_SL005 clusters with some of the Serbian isolates) or even across continents (US_JW002 clusters with Serbian and Swedish isolates).

**Figure 4.**
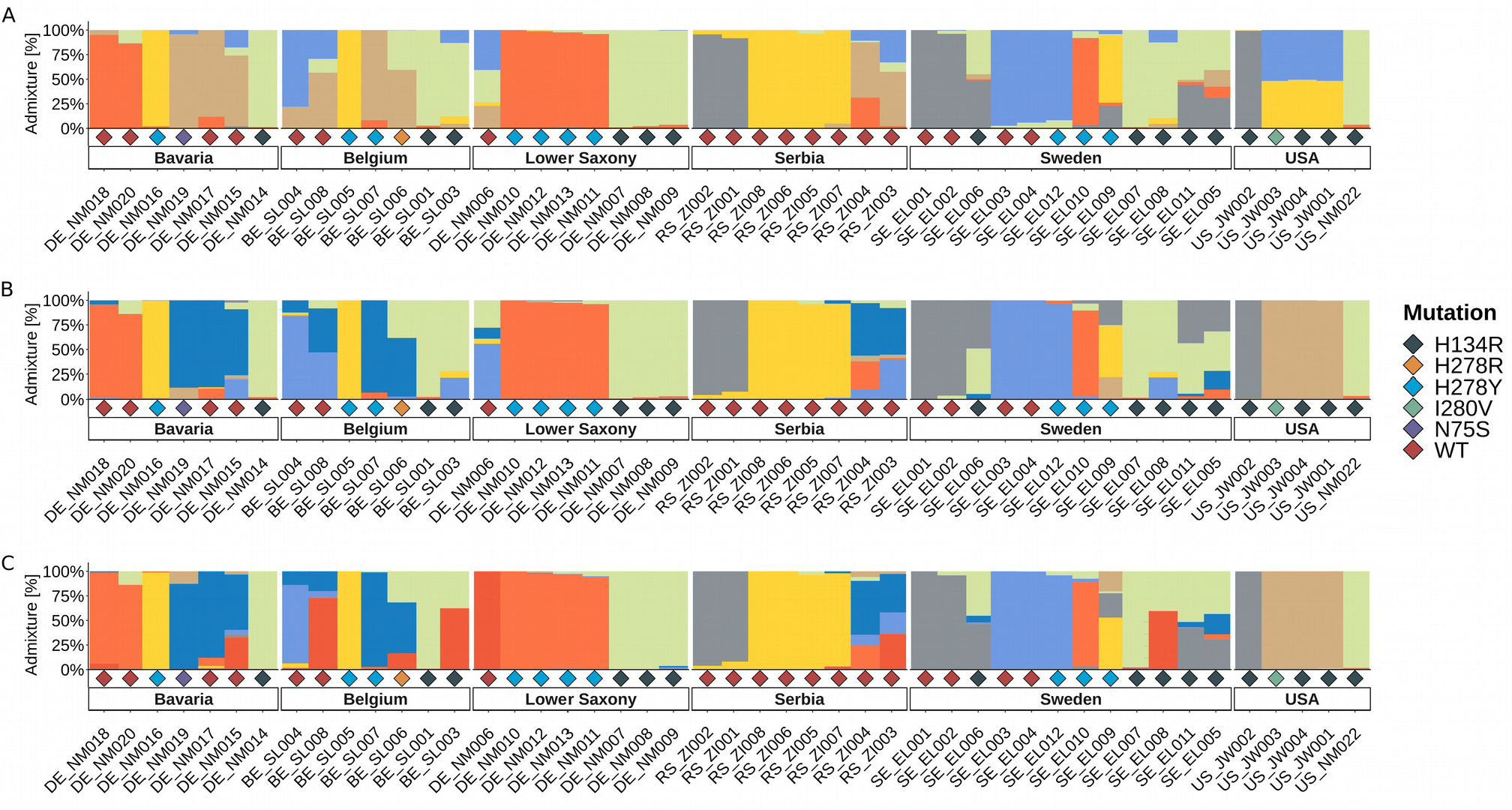
Genotypes and SDHI-mutations are not linked to sample locality and appear in different locations independently. Ancestry analysis was performed on *A. solani* samples (n=47) using the R-package LEA [*snmf(K=1:15, rep=10, ploidy=1)*] for 6 (A), 7 (B) and 8 (C) ancestral populations and then sorted according to sample locality. Colors represent individual genotypes assigned by *sparse nonnegative matrix factorization*. Bar plots show the admixture of each genotype in every sample. Diamonds represent mutations against SDHIs identified via analysis xyz.

### SDHI mutations arose in different genotypes

To understand the rise and spread of SHDI fungicide resistance, we overlaid the genotyping data with our data on SDHI-target mutations. The H134R and H278Y mutations occur 15 and 10 times in our dataset, respectively. H134R is found in every locality, except Serbia; H278Y can be found in all locations but Serbia and the US. Furthermore, we found three mutations (H278R, I280V & N75S) that occur only once in the dataset. In 19 samples, we found no mutation related to SDHI fungicides. Mutation-free isolates were collected in all different localities except for the US.

Overlaying the data shows that the mutations are not linked to the background genotypes that we defined previously. The different Sdh SNPs were found in different genetic backgrounds (Figure 3B). E.g. H134R occurs in the grey, brown and green genotype. H278Y occurs in the blue, red and yellow genotype. This strongly indicates that mutations in the SDH subunits arose independently in different genetic *A. solani* backgrounds.

## Discussion

### Diversity in context of long distance dispersal

In this study we present the first Europe-wide genetic diversity study of the early blight pathogen *Alternaria solani* using whole genome data. We used 43 isolates collected in five different localities ranging from Sweden to Serbia. Seeing that *A. solani* is considered a clonal pathogen (synonymous: mitosporic fungus, Deuteromycete, *fungi imperfecti*), we expected to see certain geographical patterns related to the genetic diversity. The strongest geographical patterns can be observed in invasive pathogens, like the rice and wheat blast fungus *Magnaporthe oryzae*, where multiple genetic lineages can be found in Southeast Asia, but single invasive lineages spread elsewhere and sometimes became epidemic (Islam *et al*., 2016; Gladieux *et al*., 2018). However, clear geographical patterns in population structure can also be observed in cosmopolitan pathogen species like the oomycete potato pathogen *Phytophthora infestans*, the causal agent of late blight, or in other ascomycete pathogens such as the sugar beet pathogen *Cercospora beticola* (Knight *et al*., 2019) and the well-studied wheat pathogen *Zymoseptoria tritici* (Vagndorf *et al*., 2018).

We did not observe any geographical clustering of *A. solani* isolates in this study. This could be an artifact of the relatively small sample size per locality. However, some pathogens do already show clear separation even when taken such small samples (Mariette *et al*., 2016). High genotype diversity and lack of isolation by distance has previously been observed for *A. solani* in an SSR-based study with populations from China. Assumed clones were detected in multiple populations separated by thousands of kilometers and random association among loci was found in half of the populations assayed (Meng et al., 2015). Our whole-genome data now suggest that these results were not an artifact of the low-resolution markers used.

For several other supposedly clonal ascomycete pathogens, such lack of isolation by distance has been seen as well. For example, in a study on the Brassica pathogen *Leptosphaeria maculans* on a regional scale similar to our sample sites in Germany (north - south France) (Travadon *et al*., 2011) or on a global scale for the barley pathogen *Ramularia collo-cygni* (Stam *et al*., 2019). *Vice versa*, isolates from distant sites cluster together on the phylogenetic tree, suggesting long distance transport e.g. by wind or seed tubers.

Adhakiri *et al*. compared field population diversity of three *Alternaria* species on potato and tomato in North Carolina and Wisconsin and found that *A. solani* had relatively lower diversity than *A. alternata* and *A. linariae*. Yet, four different haplotypes were found in one field and haplotype divergence was found in all three species. (Adhikari *et al*., 2020). Our analyses revealed up to 9 definable genotypes in *A. solani* in our sample set. Deeper analyses showed that some of the collected isolates from these genotypes are, at least likely purely clonal. They contain very few SNPs between them. Whether such clones perform worse or better under specific climatic conditions or on specific cultivars remains to be investigated.

Based on the results of our study, we conclude that the mutations in the genes coding for the different SDH subunits that lead to resistance or higher tolerance to SDHIs can be found in multiple genetic backgrounds. This indicates that the mutations occurred multiple times at different sites, rather than that a single resistant isolate emerged and spread throughout Europe. Evolution of tolerance against QoI fungicides was initially suggested to have arisen once, due to its association with an arbitrarily assigned genotype (GT I), and then spread. However, recent work showed that the F129L mutation in the cytochrome b target gene could also be observed since 2016 in another genotype (Nottensteiner *et al*., 2019). Our study further confirms the genetic heterogeneity of QoI tolerant isolates, as all of the isolates in this study show the F129L mutation. A recent study in *Z. tritici*, concluded that azole resistance is likely the result of so-called “ hotspot evolution” with convergent changes in small sets of loci and more population-specific allele frequency changes (Hartmann *et al*. 2020). Fungicides can form a major bottleneck for fungal populations, but the ability to recombine, allowed population specific adaptation in *Z. tritici* populations. Even though we found that fungicide resistance or fungicide tolerance mutations arose multiple times independently, future studies looking deeper into the effects of fungicide induced bottlenecks in *A. solani* are now required. The absence of apparent sexual recombination and the fact that true clones can be found across continents, potentially amplifies the spread of more aggressive genotypes. Bauske *et al*. (2017) found that *A. solani* isolates possessing the *SdhD*-D123E mutation were significantly more aggressive *in vivo* compared with wild-type isolates.

Fungicide pressure and the aggressiveness of *A solani* isolates under certain conditions or on a certain host cultivars alone might not be the only determinants for *A. solani* diversity in the field. Early blight lesions often appear to contain a mix of *Alternaria* species. *A. solani* often co-occurs with *A. alternata* (e.g. Zheng & Wu, 2013). Tymon *et al* also isolated several other small and large spored species from lesions on potato fields (Tymon *et al*., 2016). And also in tomato early blight symptoms can be associated with multiple species, the large spored *A. solani, A. linariae* and *A grandis*, and the small spored *A alternata* (Bessadat *et al*., 2017). Ding *et al*. found, that in some years the disease severity in the field is more strongly correlated to the presence of early inoculum *of A. alternata* (Ding *et al*., 2020). Using barcode-sequencing, they found that *A. alternata* possessed a greater diversity in the field (Ding et al., 2018). In all studies mentioned above, the disease phenotypes of *A. solani* isolates were more severe and *A. solani* is the more dominant partner in the interaction. Studies on Alternaria leaf spot on rape seed show that the also this disease is not caused just by *A. brassicae*, but that it can also be caused by *A. japonica* and that the ratio between the species and aggressiveness of the species was temperature dependent (Al-lami *et al*., 2020). Findings like these indicate that genotypic diversity of *A. solani*, can potentially also be shaped by the presence of diverse *Alternaria* species and that further research is needed to understand the effects of coinfection on early blight epidemiology.

We presented a first genome-wide diversity analysis for the early blight pathogen *A. solani* in Europe. We revealed surprising genetic diversity patterns throughout Europe and show that fungicide resistance evolved multiple times independently, rather than evolving once followed by spreading. These findings can help inform policy and fungicide management practices. The finding of multiple fungicide resistance mutation events highlights that there is “not one person to blame” for the rise of fungicide resistance in *A. solani*. This emphasizes the need to reduce the evolutionary pressure in each individual field, and keeping the population size as small as possible at all times (McDonald & Linde, 2002). New strategies using evolutionary knowledge as well as the intensive application of already non-fungicide available measures are required for good integrative disease management on a continental scale (Green *et al*., 2020). Only highly integrated approaches of pest management, including measures of all kinds, are capable of reducing the evolutionary potential of *A. solani* epidemics. To do this without increasing too much the evolutionary pressure in the field, fungicide application time frames and quantities need to be carefully adjusted as they play an important role in the development of resistances and diseases. Unfortunately, in our data-set, loss of fungicide sensitivity cannot be linked to the degree of fungicide-use, since data is not fully available for all localities in this project.

Our analyses also shows that some true clonal isolates can be found thousands of kilometers apart. Understanding *A. solani* dispersal patterns over long distances will become increasingly more relevant. Recent studies have shown clear differences in aggressiveness in different *A. solani* isolates in general, e.g. not associated with specific fungicide target mutations (Mphahlele *et al*., 2020). With the high-resolution (whole genome) genotype data presented in this study, more meaningful comparisons can be made to study the link between pathogen genotype and aggressiveness. The data will also allow for studies tracing exact clones of *A. solani* over time.

## Acknowledgements

We like to thank Regine Dittebrandt and Lena Forster for help with *A. solani* propagation.

## Supplementary Figures

**Figure S1.**
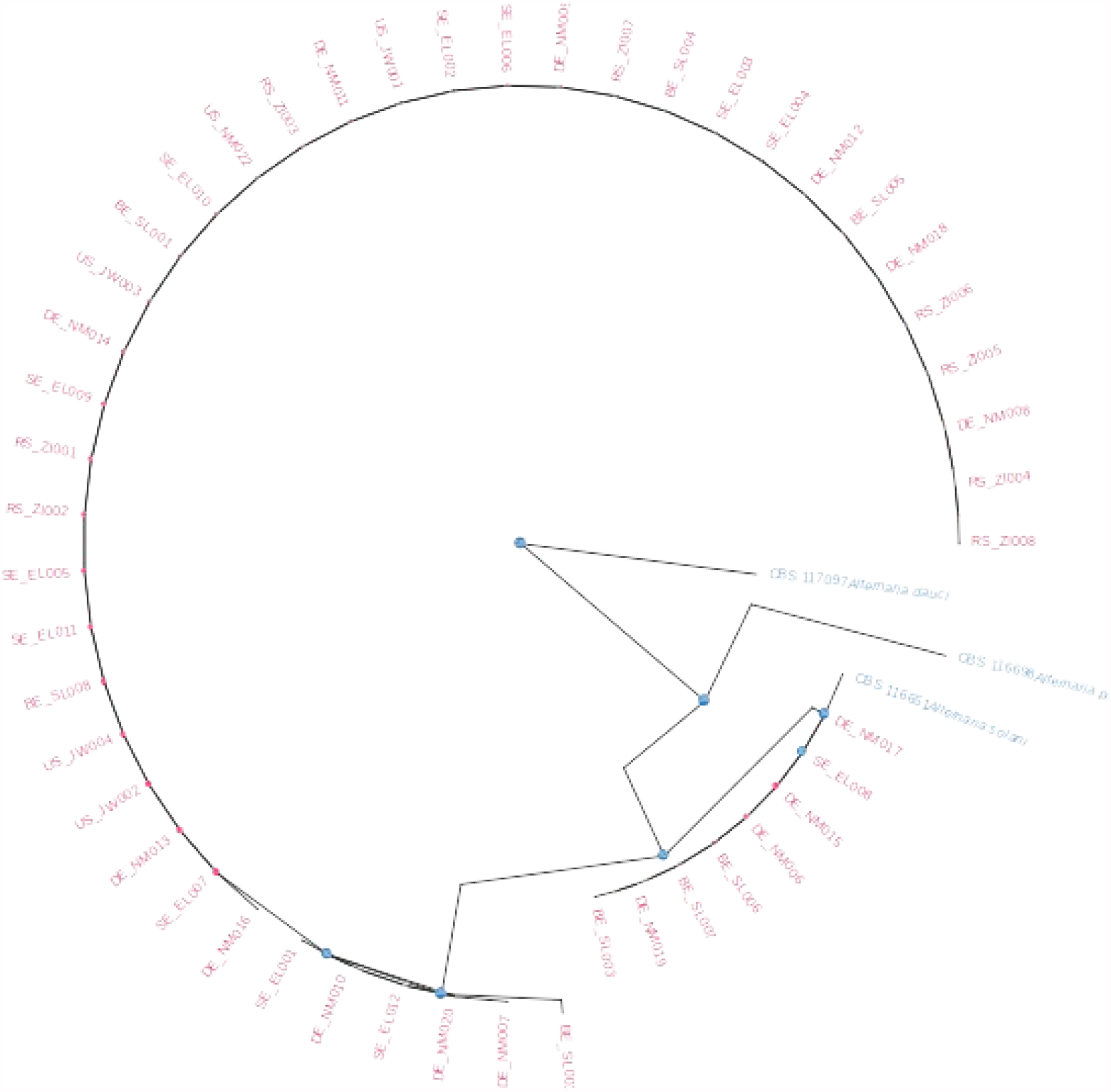

**Figure S2.**
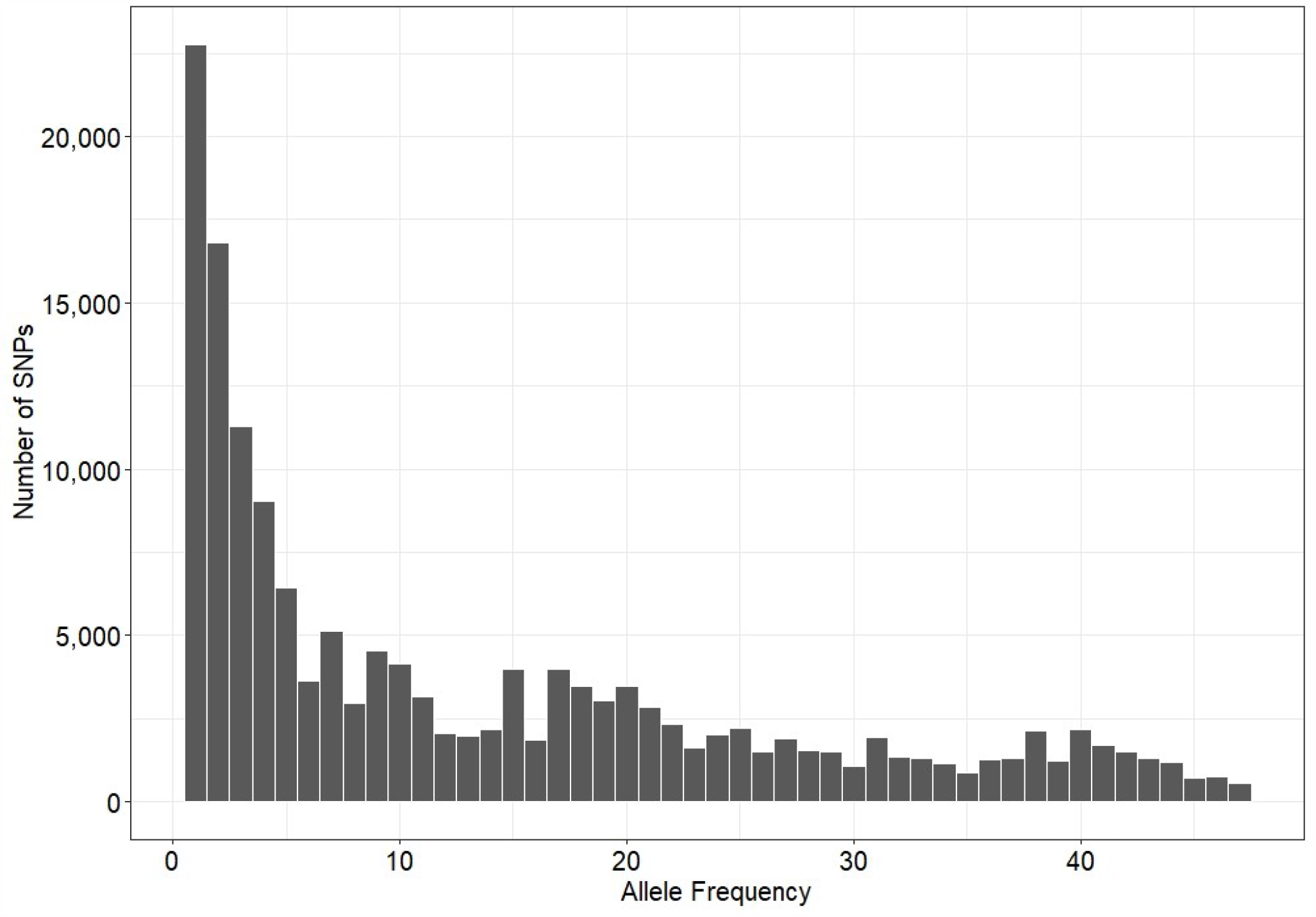

**Figure S3.**
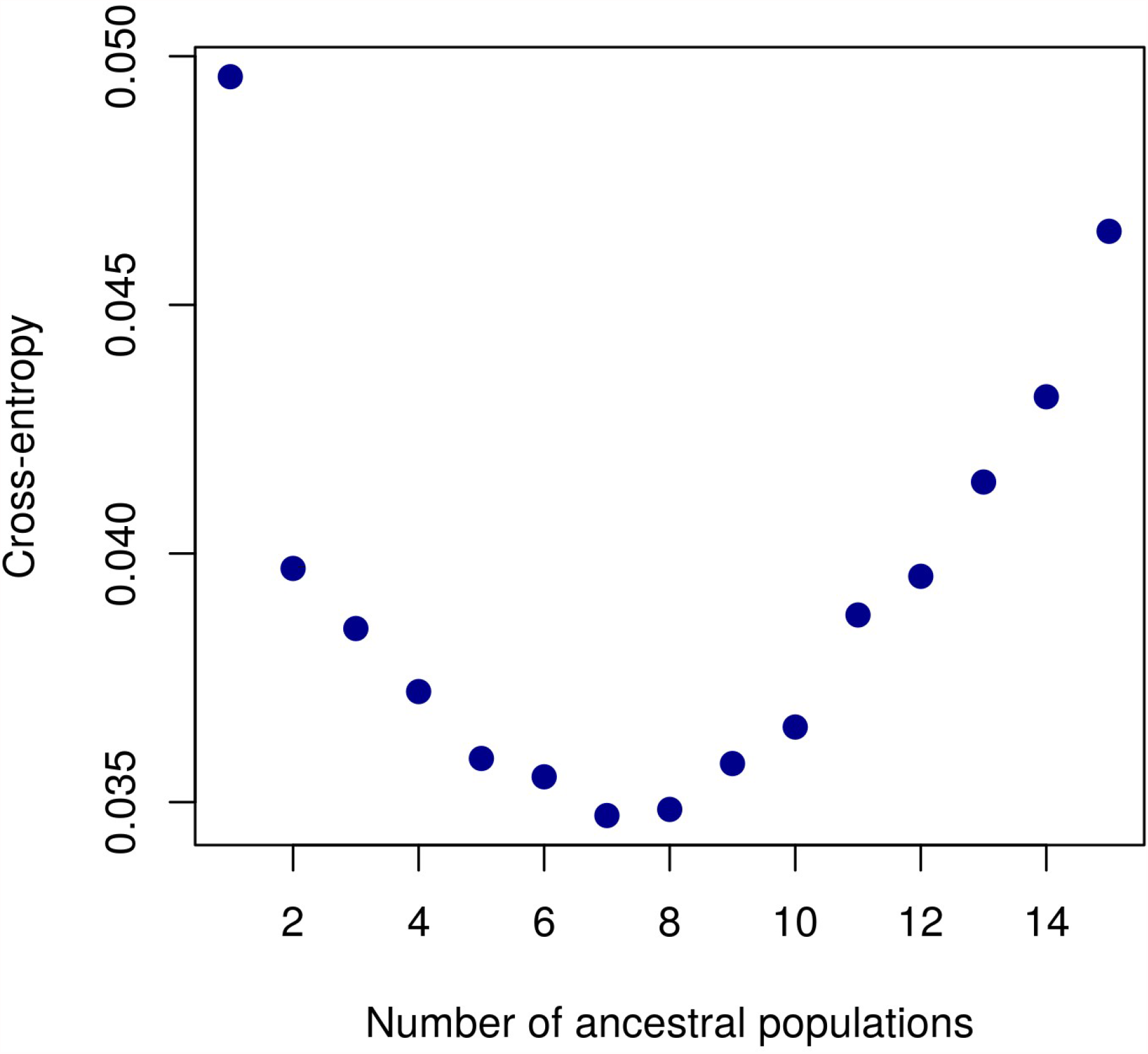

**Figure S4.**
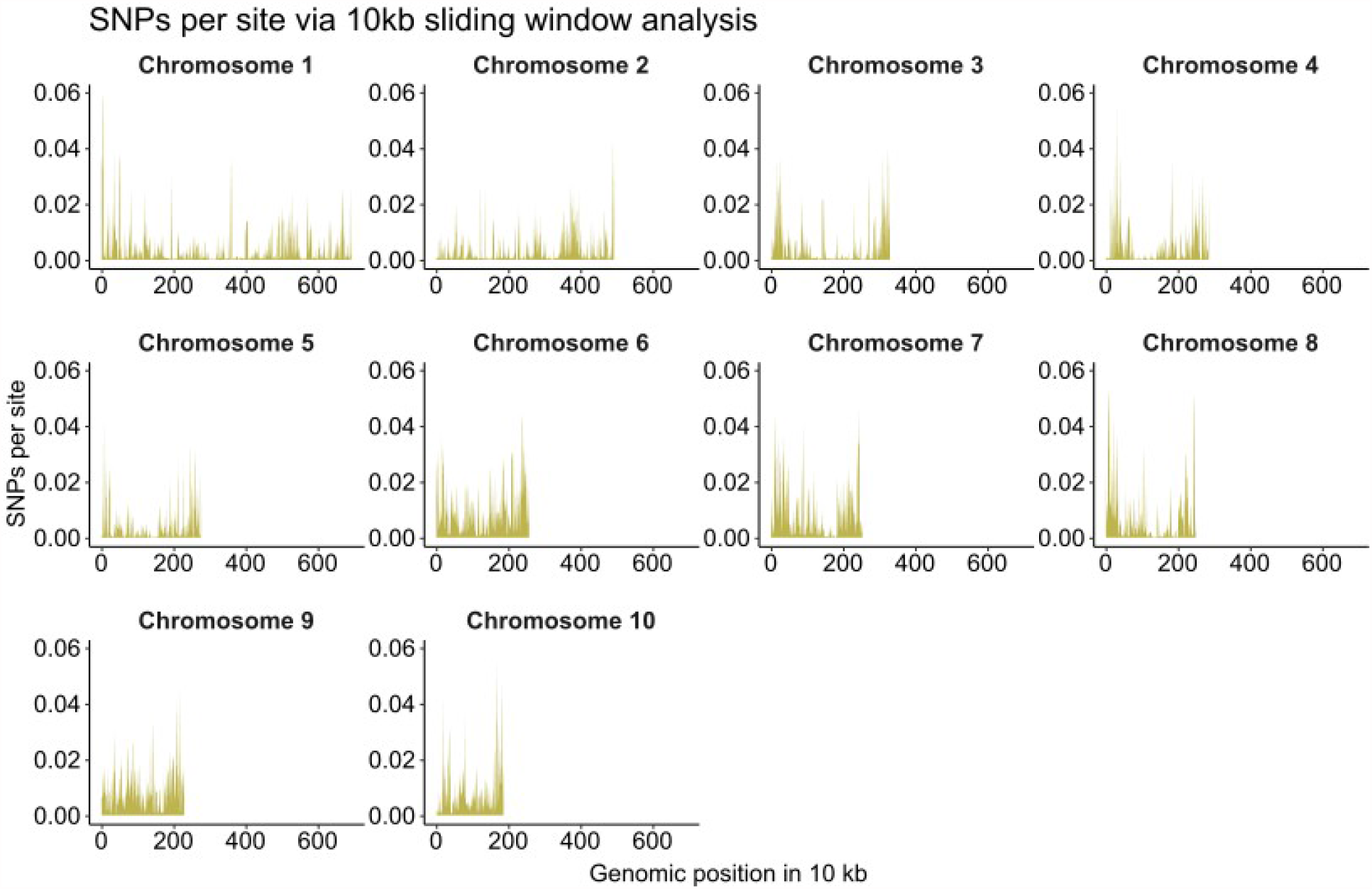

## References

Adhikari TB, Ingram T, Halterman D, Louws FJ. 2020. Gene Genealogies Reveal High Nucleotide Diversity and Admixture Haplotypes Within Three Alternaria Species Associated with Tomato and Potato. Phytopathology® 110: 1449–1464.

Adhikari TB, Knaus BJ, Grünwald NJ, Halterman D, Louws FJ. 2019. Inference of Population Genetic Structure and High Linkage Disequilibrium Among Alternaria spp. Collected from Tomato and Potato Using Genotyping by Sequencing. bioRxiv: 827790.

Al-lami HFD, You MP, Barbetti MJ. 2020. Temperature Drives Contrasting Alternaria Leaf Spot Epidemic Development in Canola and Mustard Rape from Alternaria japonica and A. brassicae. Plant Disease 104: 1668–1674.

Bartlett DW, Clough JM, Godwin JR, Hall AA, Hamer M, Parr-Dobrzanski B. 2002. The strobilurin fungicides. Pest Management Science 58: 649–662.

Bauske MJ, Mallik I, Yellareddygari SKR, Gudmestad NC. 2017. Spatial and Temporal Distribution of Mutations Conferring QoI and SDHI Resistance in Alternaria solani Across the United States. Plant Disease 102: 349–358.

Bessadat N, Berruyer R, Hamon B, Bataille-Simoneau N, Benichou S, Kihal M, Henni DE, Simoneau P. 2017. Alternaria species associated with early blight epidemics on tomato and other Solanaceae crops in northwestern Algeria. European Journal of Plant Pathology 148: 181–197.

Blake JJ, Gosling P, Fraaije BA, Burnett FJ, Knight SM, Kildea S, Paveley ND. 2018. Changes in field dose-response curves for demethylation inhibitor (DMI) and quinone outside inhibitor (QoI) fungicides against Zymoseptoria tritici, related to laboratory sensitivity phenotyping and genotyping assays. Pest Management Science 74: 302–313.

Cingolani P, Patel VM, Coon M, Nguyen T, Land SJ, Ruden DM, Lu X. 2012a. Using Drosophila melanogaster as a Model for Genotoxic Chemical Mutational Studies with a New Program, SnpSift. Frontiers in Genetics 3: 35.

Cingolani P, Platts A, Wang LL, Coon M, Nguyen T, Wang L, Land SJ, Lu X, Ruden DM. 2012b. A program for annotating and predicting the effects of single nucleotide polymorphisms, SnpEff: SNPs in the genome of Drosophila melanogaster strain w1118; iso-2; iso-3. Fly 6: 80–92.

Danecek P, Auton A, Abecasis G, Albers CA, Banks E, DePristo MA, Handsaker RE, Lunter G, Marth GT, Sherry ST. 2011. The variant call format and VCFtools. Bioinformatics 27: 2156–2158.

Delgado-Baquerizo M, Guerra CA, Cano-Díaz C, Egidi E, Wang J-T, Eisenhauer N, Singh BK, Maestre FT. 2020. The proportion of soil-borne pathogens increases with warming at the global scale. Nature Climate Change 10: 550–554.

DePristo MA, Banks E, Poplin R, Garimella KV, Maguire JR, Hartl C, Philippakis AA, del Angel G, Rivas MA, Hanna M, et al. 2011. A framework for variation discovery and genotyping using next-generation DNA sequencing data. Nat Genet 43: 491–8.

Ding S, Meinholz K, Gevens AJ. 2020. Spatiotemporal Distribution of Potato-Associated Alternaria Species in Wisconsin. Plant Disease 105: 149–155.

Edin E, Liljeroth E, Andersson B. 2019. Long term field sampling in Sweden reveals a shift in occurrence of cytochrome b genotype and amino acid substitution F129L in Alternaria solani, together with a high incidence of the G143A substitution in Alternaria alternata. European Journal of Plant Pathology 155: 627–641.

Everhart S, Gambhir N, Stam R. 2020. Population Genomics of Filamentous Plant Pathogens—A Brief Overview of Research Questions, Approaches, and Pitfalls. Phytopathology® 111: 12–22.

Frichot E, François O. 2015. LEA: an R package for landscape and ecological association studies. Methods in Ecology and Evolution 6: 925–929.

Gladieux P, Ravel S, Rieux A, Cros-Arteil S, Adreit H, Milazzo J, Thierry M, Fournier E, Terauchi R, Tharreau D, et al. 2018. Coexistence of Multiple Endemic and Pandemic Lineages of the Rice Blast Pathogen. mBio 9: e01806–17.

Green KK, Stenberg JA, Lankinen Å. 2020. Making sense of Integrated Pest Management (IPM) in the light of evolution. Evolutionary Applications 13: 1791–1805.

Hartmann FE, Vonlanthen T, Singh NK, McDonald MC, Milgate A, Croll D. The complex genomic basis of rapid convergent adaptation to pesticides across continents in a fungal plant pathogen. Molecular Ecology n/a.

Islam MT, Croll D, Gladieux P, Soanes DM, Persoons A, Bhattacharjee P, Hossain MdS, Gupta DR, Rahman MdM, Mahboob MG, et al. 2016. Emergence of wheat blast in Bangladesh was caused by a South American lineage of Magnaporthe oryzae. BMC Biology 14: 84.

Kaindl L, Small C, Stam R. 2021. AB12PHYLO: an integrated pipeline for Maximum Likelihood phylogenetic inference from ABI trace data. bioRxiv: 2021.03.01.433007.

Knaus BJ, Grünwald NJ. 2017. vcfr: a package to manipulate and visualize variant call format data in R. Molecular Ecology Resources 17: 44–53.

Knight NL, Vaghefi N, Kikkert JR, Bolton MD, Secor GA, Rivera VV, Hanson LE, Nelson SC, Pethybridge SJ. 2019. Genetic Diversity and Structure in Regional Cercospora beticola Populations from Beta vulgaris subsp. vulgaris Suggest Two Clusters of Separate Origin. Phytopathology® 109: 1280–1292.

Kozlov AM, Darriba D, Flouri T, Morel B, Stamatakis A. 2019. RAxML-NG: a fast, scalable and user-friendly tool for maximum likelihood phylogenetic inference. Bioinformatics 35: 4453–4455.

Landschoot S, Carrette J, Vandecasteele M, De Baets B, Höfte M, Audenaert K, Haesaert G. 2017. Boscalid-resistance in Alternaria alternata and Alternaria solani populations: An emerging problem in Europe. Crop Protection 92: 49–59.

Leiminger JH, Adolf B, Hausladen H. 2014. Occurrence of the F129L mutation in Alternaria solani populations in Germany in response to QoI application, and its effect on sensitivity. Plant Pathology 63: 640–650.

Leiminger JH, Auinger H-J, Wenig M, Bahnweg G, Hausladen H. 2016. Genetic variability among Alternaria solani isolates from potatoes in Southern Germany based on RAPD-profiles. Journal of Plant Diseases and Protection 120: 164–172.

Leiminger JH, Hausladen H. 2012. Early Blight Control in Potato Using Disease-Orientated Threshold Values. Plant Disease 96: 124–130.

Leiminger JH, Hausladen H. 2014. Untersuchungen zur Befallsentwicklung und Ertragswirkung der Dürrfleckenkrankheit (Alternaria spp.) in Kartoffelsorten unterschiedlicher Reifegruppe. Gesunde Pflanzen 66: 29–36.

Li H, Durbin R. 2009. Fast and accurate short read alignment with Burrows-Wheeler transform. Bioinformatics (Oxford, England) 25: 1754–1760.

Mallik I, Arabiat S, Pasche JS, Bolton MD, Patel JS, Gudmestad NC. 2014. Molecular characterization and detection of mutations associated with resistance to succinate dehydrogenase-inhibiting fungicides in Alternaria solani. Phytopathology 104: 40–49.

Mariette N, Androdias A, Mabon R, Corbière R, Marquer B, Montarry J, Andrivon D. 2016. Local adaptation to temperature in populations and clonal lineages of the Irish potato famine pathogen Phytophthora infestans. Ecology and Evolution 6: 6320–6331.

McDonald BA, Linde C. 2002. Pathogen population genetics, evolutionary potential, and durable resistance. Annu Rev Phytopathol 40: 349–79.

McKenna A, Hanna M, Banks E, Sivachenko A, Cibulskis K, Kernytsky A, Garimella K, Altshuler D, Gabriel S, Daly M, et al. 2010. The Genome Analysis Toolkit: A MapReduce framework for analyzing next-generation DNA sequencing data. Genome Research 20: 1297–1303.

Metz N, Adolf B, Chaluppa N, Hückelhoven R, Hausladen H. 2019. Occurrence of sdh Mutations in German Alternaria solani Isolates and Potential Impact on Boscalid Sensitivity In Vitro, in the Greenhouse, and in the Field. Plant Disease 103: 3065–3071.

Mphahlele GH, Kena MA, Manyevere A. 2020. Evaluation of aggressiveness of Alternaria solani isolates to commercial tomato cultivars. Archives of Phytopathology and Plant Protection 53: 570–580.

Nottensteiner M, Absmeier C, Zellner M. 2019. QoI Fungicide Resistance Mutations in Alternaria solani and Alternaria alternata are Fully Established in Potato Growing Areas in Bavaria and Dual Resistance against SDHI Fungicides is Upcoming. Gesunde Pflanzen 71: 155–164.

Odilbekov F, Edin E, Garkava-Gustavsson L, Hovmalm HP, Liljeroth E. 2016. Genetic diversity and occurrence of the F129L substitutions among isolates of Alternaria solani in south-eastern Sweden. Hereditas 153: 10.

Odilbekov F, Edin E, Mostafanezhad H, Coolman H, Grenville-Briggs LJ, Liljeroth E. 2019. Within-season changes in Alternaria solani populations in potato in response to fungicide application strategies. European Journal of Plant Pathology 155: 953–965.

Pereira DA, McDonald BA, Brunner PC. 2017. Mutations in the CYP51 gene reduce DMI sensitivity in Parastagonospora nodorum populations in Europe and China. Pest Management Science 73: 1503–1510.

Pfeifer B, Wittelsbürger U, Ramos-Onsins SE, Lercher MJ. 2014. PopGenome: an efficient Swiss army knife for population genomic analyses in R. Molecular Biology and Evolution 31: 1929–1936.

Rotem J. 1994. The genus Alternaria: biology, epidemiology, and pathogenicity. St Paul, USA: American Phytopathological Society.

Stam R, Sghyer H, Tellier A, Hess M, Hückelhoven R. 2019. The Current Epidemic of the Barley Pathogen Ramularia collo-cygni Derives from a Population Expansion and Shows Global Admixture. Phytopathology® 109: 2161–2168.

Stamatakis A. 2014. RAxML version 8: a tool for phylogenetic analysis and post-analysis of large phylogenies. Bioinformatics 30: 1312–1313.

Travadon R, Sache I, Dutech C, Stachowiak A, Marquer B, Bousset L. 2011. Absence of isolation by distance patterns at the regional scale in the fungal plant pathogen Leptosphaeria maculans. Fungal Biology 115: 649–659.

Tymon LS, Peever TL, Johnson DA. 2016. Identification and Enumeration of Small-Spored Alternaria Species Associated with Potato in the U.S. Northwest. Plant Disease 100: 465–472.

Upadhyay P, Ganaie SH, Singh N. 2019. Diversity Assessment Among Alternaria solani Isolates Causing Early Blight of Tomato in India. Proceedings of the National Academy of Sciences, India Section B: Biological Sciences 89: 987–997.

Vagndorf N, Heick TM, Justesen AF, Andersen JR, Jahoor A, Jørgensen LN, Orabi J. 2018. Population structure and frequency differences of CYP51 mutations in Zymoseptoria tritici populations in the Nordic and Baltic regions. European Journal of Plant Pathology 152: 327–341.

van der Waals JE, Korsten L, Slippers B. 2004. Genetic Diversity Among Alternaria solani Isolates from Potatoes in South Africa. Plant Disease 88: 959–964.

Wang L-G, Lam TT-Y, Xu S, Dai Z, Zhou L, Feng T, Guo P, Dunn CW, Jones BR, Bradley T, et al. 2020. Treeio: An R Package for Phylogenetic Tree Input and Output with Richly Annotated and Associated Data. Molecular Biology and Evolution 37: 599–603.

Weber B, Halterman DA. 2012. Analysis of genetic and pathogenic variation of Alternaria solani from a potato production region. European Journal of Plant Pathology 134: 847–858.

Wickham H. 2009. ggplot2: Elegant Graphics for Data Analysis. Springer-Verlag New York.

Wolters PJ, Faino L, van den Bosch TBM, Evenhuis B, Visser RGF, Seidl MF, Vleeshouwers VGAA. 2018. Gapless Genome Assembly of the Potato and Tomato Early Blight Pathogen Alternaria solani. Molecular plant-microbe interactions: MPMI 31: 692–694.

Woudenberg JHC, Groenewald JZ, Binder M, Crous PW. 2013. Alternaria redefined. Studies in Mycology 75: 171–212.

Zhang Y, Zhou Q, Tian P, Li Y, Duan G, Li D, Zhan J, Chen F. 2020. Induced expression of CYP51 associated with difenoconazole resistance in the pathogenic Alternaria sect. on potato in China. Pest Management Science 76: 1751–1760.

Zheng X, Levine D, Shen J, Gogarten SM, Laurie C, Weir BS. 2012. A high-performance computing toolset for relatedness and principal component analysis of SNP data. Bioinformatics (Oxford, England) 28: 3326–3328.

Zheng HH, Wu XH. 2013. First Report of Alternaria Blight of Potato Caused by Alternaria tenuissima in China. Plant Disease 97: 1246–1246.

Slowikowski, K. 2018. ggrepel: Automatically position non-overlapping text labels with ‘ggplot2.’ R Package Version 0.8. 0 Ed.

